# Inter-organ growth coordination is mediated by the Xrp1/Dilp8 axis in *Drosophila*

**DOI:** 10.1101/462473

**Authors:** Laura Boulan, Ditte Andersen, Julien Colombani, Emilie Boone, Pierre Léopold

**Affiliations:** Institut Curie, PSL Research University, CNRS UMR3215, INSERM U934, UPMC Paris-Sorbonne, 26 Rue d’Ulm, 75005, Paris, France; Université Côte d’Azur, CNRS UMR7277, Inserm U1091, iBV, Parc Valrose, 06108, Nice, France

**Keywords:** organ growth, Dilp8, Drosophila, Minute, Xrp1, Coordination, RpS12

## Abstract

How organs scale with other body parts is not mechanistically understood. We have addressed this question using the *Drosophila* imaginal disc model. When growth of one disc domain is perturbed, other parts of the disc and other discs slow down their growth, maintaining proper inter-disc and intra-disc proportions. We show here that the relaxin-like Dilp8 is required for this inter-organ coordination. Our work also reveals that the stress-response transcription factor Xrp1 plays a key role upstream of *dilp8* in linking organ growth status with non-autonomous/systemic growth response. In addition, we show that the small ribosomal subunit protein RpS12 is required to trigger Xrp1-dependent non-autonomous response. Our work demonstrates that RpS12, Xrp1 and Dilp8 constitute a new, independent regulatory module that ensures intra- and inter-organ growth coordination during development.

## INTRODUCTION

Body size and organ proportions are important characteristics that define animal fitness, mobility, predation and competition for a given species. Animals adapt their growth and body size to environmental conditions, but the relative proportions between the different body parts remain remarkably constant, suggesting that coordination mechanisms are at play during development. Most recent studies on growth have focused on the mechanism of organ size determination without interrogating the complex question of coordinating growth between organs. One elegant way to tackle this question is to induce a local growth perturbation in a given organ and to analyze the non-autonomous/systemic responses triggered by this perturbation. This approach was recently used in mice where unilateral inhibition of proliferation in the limb cartilage reduces contralateral bone growth, thereby contributing to the maintenance of left/right bone symmetry (Roselló-Díez et al., 2018). In *Drosophila*, growth inhibition in the wing imaginal disc during larval development triggers a non-autonomous response reducing the growth rate of the leg or eye disc. As a consequence, the relative proportions between the slow-growing tissue and unperturbed tissues are maintained throughout development (Jaszczak and Halme, 2016;Parker and Shingleton, 2011). Therefore, it seems that conserved mechanisms allow slow-growing tissues to systemically act on general growth parameters and maintain proportions between body parts. What signals mediate this inter-organ communication, and what molecular mechanisms link growth inhibition and signal production are currently open questions.

Here we show that the relaxin-like Dilp8 (Colombani et al., 2012;Garelli et al., 2012) is the signal that triggers the coordination of growth among organs during *Drosophila* development. Furthermore, we identify a novel pathway relying on the transcription factor Xrp1, which triggers Dilp8 expression in slow-growing tissues, allowing inter-organ coordination. The JNK and Hippo signaling pathways, both previously shown to regulate Dilp8, are not involved in this process. Our results also indicate that the small ribosomal subunit protein RpS12 is required to trigger the Xrp1-dependent non-autonomous response.

## RESULTS

### Dilp8 is required to coordinate growth of imaginal tissues

Imaginal discs grow extensively during the larval phase to form adult body parts after metamorphosis. In the “wing” imaginal disc, the pouch gives rise to the adult wing blade whereas the hinge and notum form the proximal and thorax structures, respectively (Figure 1A). As an experimental setup to induce local growth perturbation in larval discs, we used the *pdm2-Gal4* line to drive expression specifically in the wing pouch, combined with an RNAi line targeting a ribosomal protein (*UAS-RpL7^RNAi^*). As expected, *pdm2>RpL7^RNAi^* animals have smaller wing pouch territories at 4 days (90 hours AED) and 5 days (110 hours AED) of development compared to controls (“minute” discs, Figure 1B,D). In these conditions, we also observed a non-autonomous inhibition of growth in the eye discs, indicative of inter-organ coordination (Figure 1G). Notably, the hinge and the notum, two independent territories of the wing imaginal disc, also display coordinated growth reduction (Figure 1C,D). To quantify these non-autonomous responses, hereafter referred to as inter/intra-organ coordination (IOC), we chose to evaluate the ratio between hinge+notum and pouch surfaces in our following experiments.

**Figure 1.**
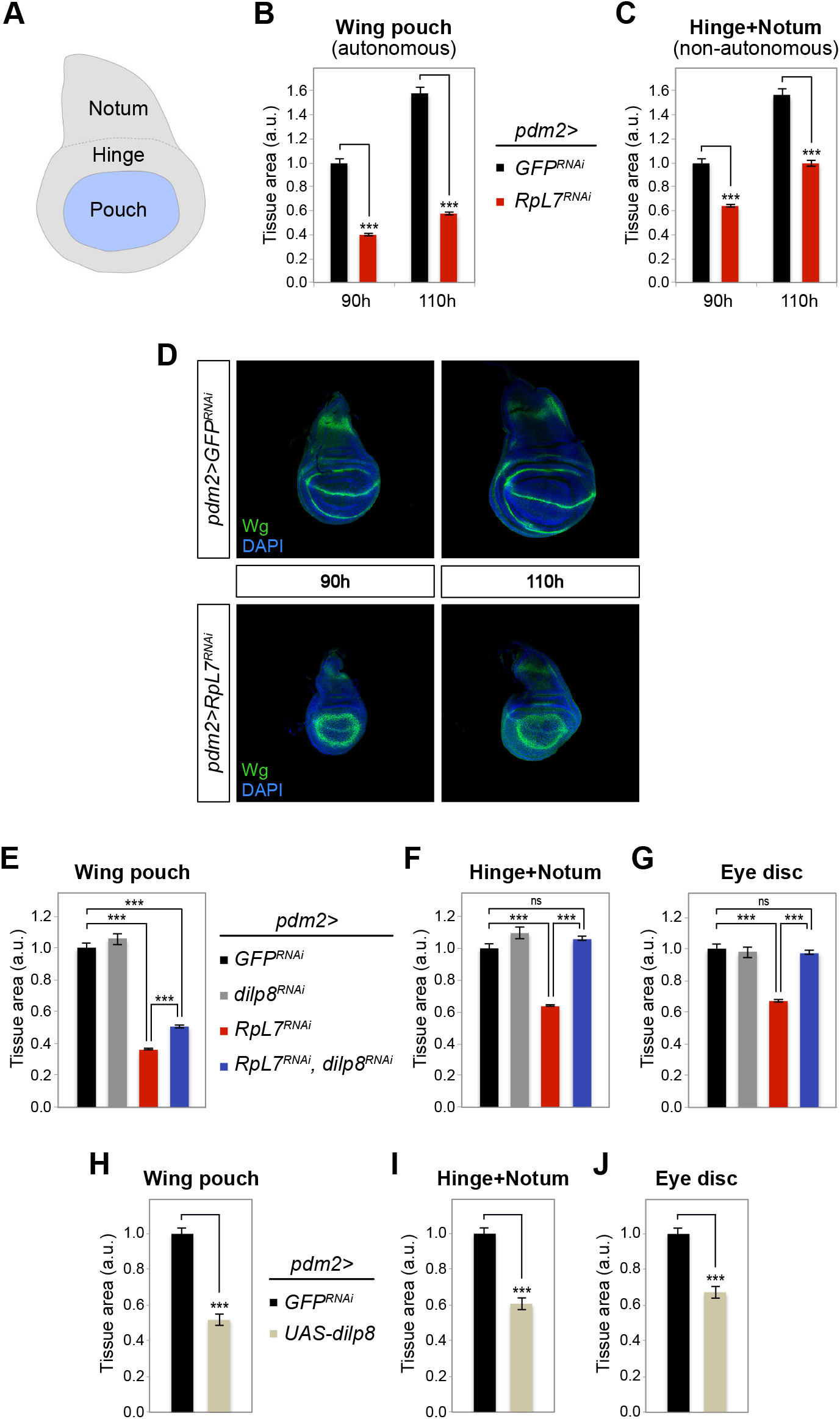
Dilp8 mediates growth coordination in response to local growth perturbations. (A) Scheme of a wing imaginal disc showing the different territories used to assess tissue growth. (B,C) Area measurements of the wing pouch (autonomous; B) and hinge and notum (non-autonomous; C) in control wing discs (*pdm>GFP^RNAi^*) and minute wing discs (*pdm2>RpL7^RNAi^*), showing the coordination of growth between slow-growing tissues and other imaginal tissues at 90h and 110h AED (n≥29). (D) Representative pictures of wing imaginal discs of the two genotypes stained for Wingless (Wg), illustrating the coordinated reduction in area size of wing and notum/hinge in *pdm2>RpL7^RNAi^* animals. (E-J) Area measurement of the wing pouch (E and H), the adjacent disc territories (hinge and notum; F and I) and a remote organ (eye disc; G and J) in animals of the different genotypes. Experiments were done at 110h AED. (E-G) The downregulation of *dilp8* in *pdm2>RpL7^RNAi^* animals (*pdm2>RpL7^RNAi^, dilp8^RNAi^*) has a very mild effect on the size of the wing pouch but fully rescues growth non-autonomously (n≥27). (H-J) *dilp8* overexpression in the wing pouch (*pdm2>dilp8*) triggers both autonomous and non-autonomous inhibition of tissue growth (n≥26). Data are represented as mean ± SEM (^***^ p<0.001 and ns=not significant, ANOVA).

Since Dilp8 is (i) highly secreted in conditions of growth impairment and (ii) acting as an inhibitor of ecdysone production and tissue growth, we tested its role in IOC. RNAi-mediated inhibition of *dilp8* in the slow-growing wing pouch (*pdm2>rpl7^RNAi^, dilp8^RNAi^*) fully rescued growth in the hinge and notum territories but not in the pouch (Figure 1E,F). This was confirmed using other genetic tools inducing similar growth perturbation *(elav-GAL80, nub>RpS3^RNAi^*; Figure S1A-C). Notably, Dilp8 inhibition in the pouch also rescued growth of the eye disc (Figure 1G), indicating that Dilp8 is required for growth coordination between remote organs. In addition, overexpressing Dilp8 in the wing pouch is sufficient to reduce the size of the hinge and notum territories as well as the eye disc at the same developmental time (Figure 1H-J). This overall indicates that Dilp8 act as a growth inhibitor and is a major actor of IOC.

### Dilp8-mediated IOC is independent of JNK and Hippo signaling

What is the molecular mechanism linking growth perturbation to *dilp8* upregulation and the systemic growth response? The JNK pathway activates *dilp8* expression in the context of neoplastic growth (Colombani et al., 2012), or regenerative growth through JAK/STAT signaling (Katsuyama et al., 2015), while the Hippo pathway controls *dilp8* expression in physiological conditions to buffer growth-associated developmental noise (Boone et al., 2016). We found that inhibition of JNK or Hippo signaling was not able to rescue the upregulation of *dilp8* mRNA levels observed at 110 hours AED (*elav-GAL80, nub>RpS3^RNAi^*) (Figure 2A). We noticed however a partial rescue of the developmental delay at pupariation by JNK and Hippo inhibition, suggesting that both signaling pathways could play a role after 110 hours AED to time pupariation (Figure 2B). Finally, in these conditions, inhibition of JNK or Hippo signaling was not able to rescue tissue growth, neither autonomously (wing pouch) nor non-autonomously (hinge + notum) (Figure 2C,D).

**Figure 2.**
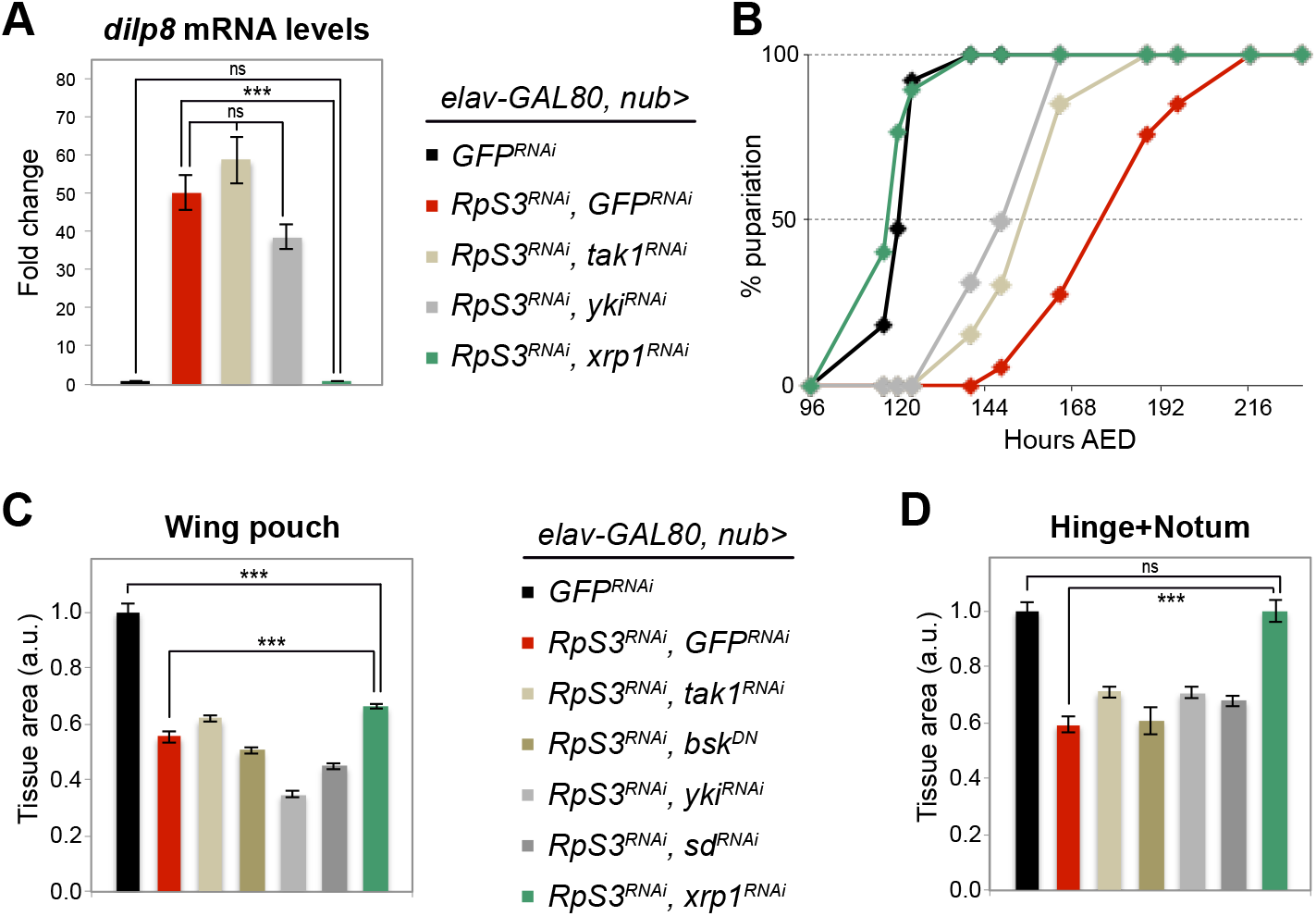
Removal of *xrp1* in minute discs reduces *dilp8* levels and prevents growth coordination. (A) Measurement of *dilp8* mRNA levels by qRT-PCR on whole larvae of the indicated genotypes. (B) Pupariation curve showing a full rescue of the delay at pupariation upon knock-down of *xrp1* in the pouch. Percentage of larvae that have pupariated at the indicated hours AED is shown (n≥53). (C,D) Measurement of pouch area (C) and hinge and notum area (D) in wing discs of the indicated genotypes, showing a rescue of the hinge+notum territories upon knock-down of *xrp1* in minute wing pouch. Inhibiting JNK or Hpo signaling does not rescue tissue growth (n≥13). (A, C, D) Experiments were done at 110h AED. Data are represented as mean ± SEM (^***^ p<0.001 and ns=not significant, ANOVA).

Additional known regulators of *dilp8* transcription include EcR (Zhang et al., 2015) and PERK/ATF4 signaling in response to ER stress (Demay et al., 2014). However, silencing these pathways in minute discs does not rescue non-autonomous growth inhibition (Figure S2). Therefore, none of the already known regulatory pathways triggering *dilp8* expression acts in condition of local growth perturbation.

### Xrp1 controls IOC upstream of Dilp8

Dilp8 was identified in a genome-wide genetic screen for genes involved in the developmental delay induced by growth impairment in the imaginal discs (Colombani et al., 2012). Another gene came out of this initial screen called *xrp1*, encoding a bZIP transcription factor. Although poorly rescuing the delay induced by neoplastic growth (*elav-GAL80, rn>avl^RNAi^*, data not shown), silencing *xrp1* very efficiently rescued the developmental delay induced by slow-growing discs (*elav-GAL80, nub>Rps3^RNAi^*; Figure 2B), and, accordingly, the levels of *dilp8* mRNA (Figure 2A). Consistent with the rescue of *dilp8* expression levels, we observed full non-autonomous rescue of tissue growth in *elav-GAL80, nub>RpS3^RNAi^, xrp1^RNAi^* animals (hinge+notum, Figure 2D), with no effect on autonomous (pouch) growth (Figure 2C). Taken together, these results indicate that the bZIP transcription factor Xrp1 is required in slow-growing tissues to upregulate Dilp8, which in turn triggers developmental delay and growth coordination.

### Xrp1 overexpression is sufficient to trigger non-autonomous reduction of tissue growth

Xrp1 has previously been implicated in the response to genotoxic stress (irradiation), the maintenance of genome integrity downstream of p53 (Akdemir et al., 2007) and during P-element dysgenesis (Francis et al., 2016). In order to gain insights into its putative growth-related function, we generated transgenic animals expressing the short and the long isoforms of Xrp1 under *UAS* control (*UAS-xrp1-S* and *UAS-xrp1-L*). *xrp1* overexpression in the wing pouch with either of the two transgenes leads to a strong autonomous reduction of tissue size (*pdm2>xrp1-S* and *pdm2>xrp1-L*, Figure 3A), accompanied by an induction of apoptosis (Figure S3A). Furthermore, non-autonomous growth inhibition was observed in the neighboring territories - hinge and notum - as well as in the eye disc (Figure 3B,C). Accordingly, *dilp8* mRNA levels were upregulated in *pdm2>xrp1-L* animals (Figure 3D). As observed in slow-growing (Rp^RNAi^) discs, concomitant downregulation of *dilp8* in *pdm2>xrp1-L* wing pouches had no effect autonomously but fully rescued growth non-autonomously (Figure 3E,F). This condition did not rescue apoptosis in the pouch (Figure S3A), suggesting that Dilp8 only mediates the non-autonomous effects of Xrp1. These results indicate that Xrp1 is sufficient to induce IOC and confirm that Dilp8 is required downstream of Xrp1 in this process.

**Figure 3.**
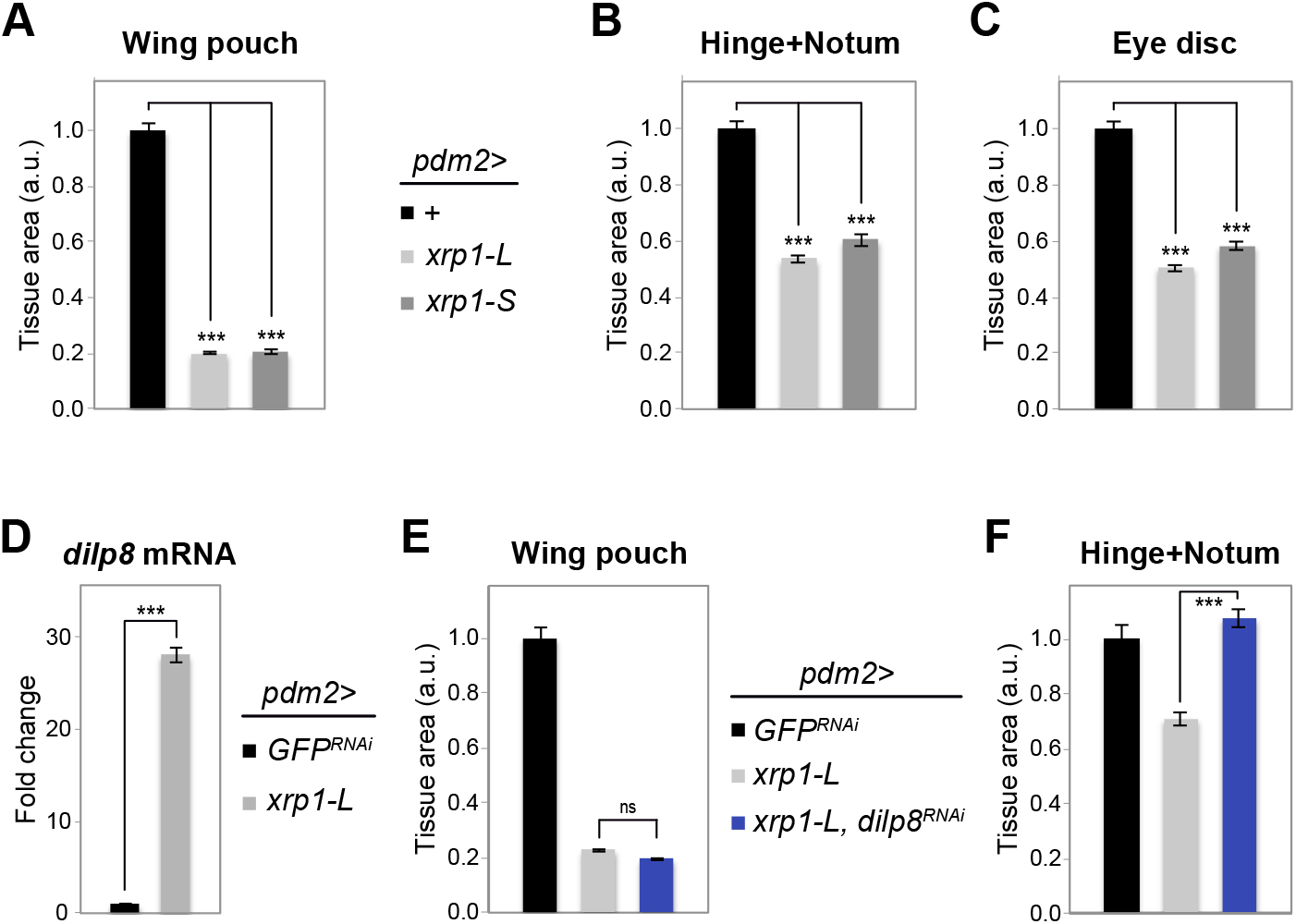
Non-autonomous growth inhibition by Xrp1 is mediated by Dilp8. (A-C) Measurement of wing pouch area (A), hinge and notum area (B) and area of a remote organ (eye disc; C) in animals of the indicated genotypes. Overexpression of the short (*UAS-xrp1-S*) and long (*UAS-xrp1-L*) isoforms of *xrp1* in the wing pouch inhibits tissue growth and triggers inter-organ growth coordination (n≥16). (D) *xrp1* overexpression in the wing pouch upregulates *dilp8* mRNA levels, as measured by qRT-PCR on whole larvae. (E, F) Measurement of wing pouch area (E), hinge and notum area (F) in wing discs of the indicated genotypes, showing that Dilp8 is required for the non-autonomous, but not the autonomous, effects of *xrp1* overexpression (n≥15). (A-F) All experiments were done at 110h AED. Data are represented as mean ± SEM (^***^ p<0.001 and ns=not significant, ANOVA).

Finally, in order to better understand the link between slow-growing discs and Xrp1 function, we analyzed *xrp1* expression in response to local growth perturbation. We observed a modest but consistent upregulation of an *xrp1-lacZ* reporter construct in slow-growing wing pouches (Figure S3B, compare Dilp8 accumulation inside and outside of the pouch domain). This result was confirmed by qRT-PCR on dissected wing imaginal discs from *elav-GAL80, nub>RpS3^RNAi^* and *>RpL7^RNAi^* animals, with primers detecting either a common region to the long and the short isoforms, or only the long one (Figure S3C). These results suggest either that a fine-tuned transcriptional regulation of Xrp1 is sufficient to trigger *dilp8* expression and IOC, or that Xrp1 is regulated at post-transcriptional levels.

### A role for the unusual ribosomal protein RpS12 in IOC

What could be the upstream regulator of Xrp1 in slow-growing tissues? Xrp1 was recently found in a screen for genes involved in minute-induced cell competition (Lee et al., 2016). Minute cells are heterozygous mutant for ribosomal protein genes, a context presenting some similarities with the RNAi-mediated downregulation of *RpL7* and *RpS3* that we used to inhibit growth in the wing discs. The only upstream regulator of Xrp1 known so far is p53, which is required to induce *xrp1* upon irradiation (Akdemir et al., 2007). In addition, previous work has shown that *xrp1* is slightly induced upon oxidative stress (Gruenewald et al., 2009). Interestingly, oxidative stress and Nrf2 are also implicated in minute cell competition (Kucinski et al., 2017). However, RNAi-mediated downregulation of *p53* or inhibition of oxidative stress signaling (*UAS-CAT+SOD*, *UAS-nrf2^RNAi^*) in slow-growing discs did not rescue non-autonomous growth as observed with *UAS-xrp1^RNAi^* (Figure S2), pointing to another mode of Xrp1 regulation in this context.

Another potential candidate for this regulation is the ribosomal protein RpS12, since it plays a role in triggering minute-induced cell competition (Kale et al., 2018). Consistent with its function as a ribosomal protein, silencing RpS12 in the wing pouch (*pdm2>rpS12^RNAi^*) led to a strong autonomous inhibition of tissue growth (Figure 4A). Yet, in contrast to the downregulation of RpS3 and RpL7, RpS12 silencing does not induce IOC (Figure 4B,C), suggesting that it is itself required for this process. We therefore tested the requirement for RpS12 in IOC induced by the knock-down of other *Rp* genes. Intriguingly, the downregulation of RpS12 in *pdm2>RpL7^RNAi^* animals had no autonomous effect in the pouch itself but rescued non-autonomous growth inhibition in the notum and hinge and well as in the eye disc (Figure 4A-C). Similar results were obtained in *elav-GAL80, nub>RpS3^RNAi^, rpS12^RNAi^* animals (Figure S4). These observations indicate that RpS12 itself is required to trigger IOC upon the reduction of ribosomal protein expression. Consistent with Dilp8 being the secreted signal required for this coordination, very little upregulation of *dilp8* was observed in *pdm2>rps12^RNAi^* discs, and the up-regulation of *dilp8* observed in the wing pouch of *pdm2>RpL7^RNAi^* animals was rescued by silencing *rpS12* (Figure 4D). Collectively, these results suggest that Rps12 functions as a sensor of ribosomal protein function upstream of Xrp1 and Dilp8 For the control of IOC.

**Figure 4.**
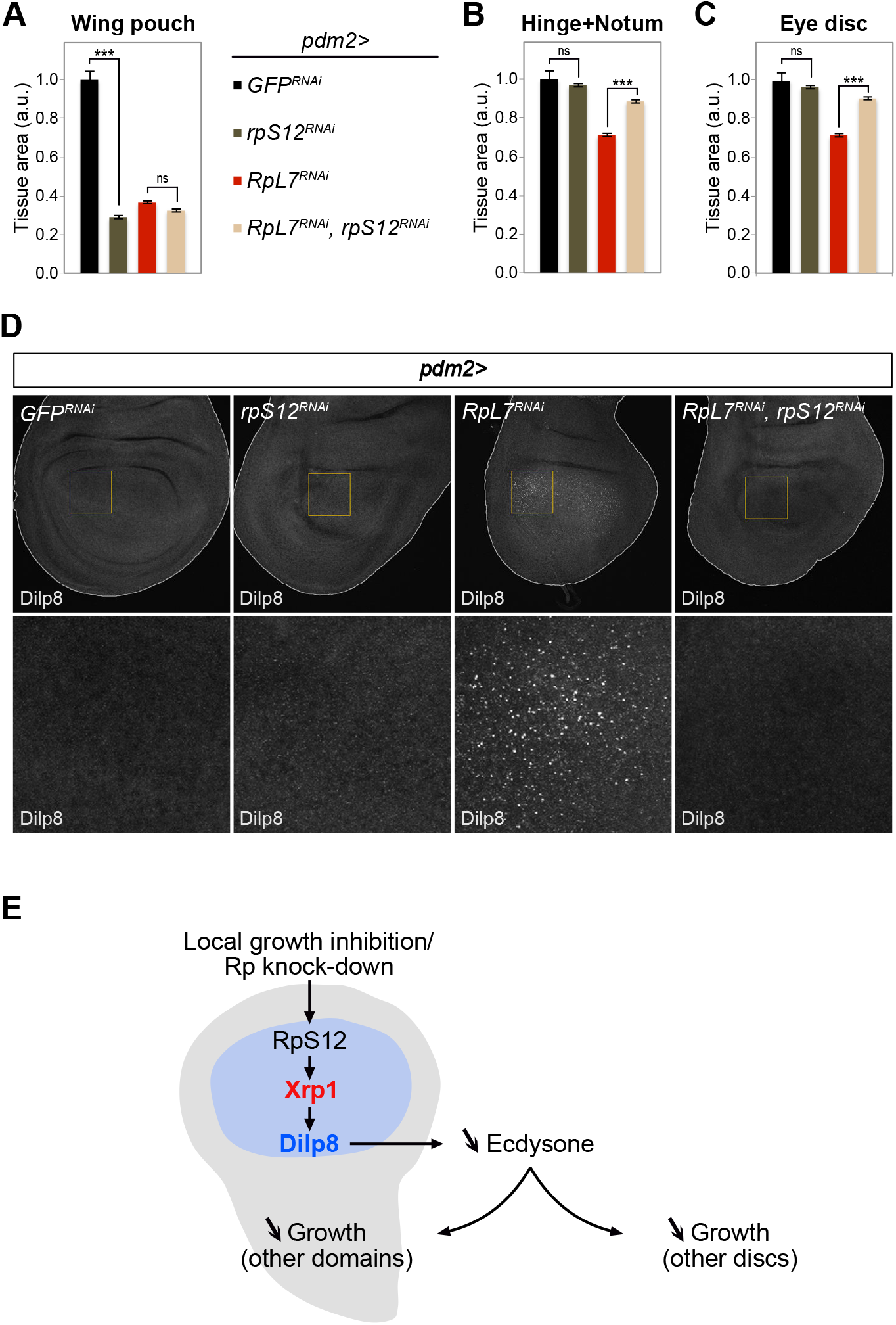
RpS12 is required for *dilp8* induction and systemic growth inhibition. (A-C) Measurement of wing pouch area (A), hinge and notum area (B) and area of a remote organ (eye disc; C) in animals of the indicated genotypes, showing that the knock-down of RpS12 in minute wing pouch rescues tissue growth in a non-autonomous manner (n≥21). The downregulation of *rpS12* alone strongly reduces tissue growth autonomously (A) but does not trigger inter-organ coordination. Data are represented as mean ± SEM (^***^ p<0.001 and ns=not significant, ANOVA). (D) Dilp8 staining in wing imaginal discs of the indicated genotypes, showing the reduction of Dilp8 levels in *pdm2>RpL7^RNAi^, rpS12^RNAi^* discs. (A-D) All experiments were done at 110h AED. (E) A model for the RpS12/Xrp1/Dilp8 regulatory pathway triggering IOC in response to local growth perturbation.

## DISCUSSION

### Dilp8 is a master regulator of systemic growth

Dilp8 is secreted by growing, regenerating and tumorous tissues. It was shown to act remotely on steroid hormone production to inhibit the transition to the pupal stage and to delay development. In this work, we show that the induction of *dilp8* in slow-growing minute wing discs is required and sufficient to trigger growth inhibition in other disc domains and other discs, thereby contributing to maintaining proper body proportions. These results establish that Dilp8 is a key regulator of the systemic response to a local growth perturbation, acting both by delaying the developmental transition and by coordinating the growth rates between organs.

Several lines of evidence indicate that Dilp8-mediated IOC requires the modulation of ecdysone levels. First, Dilp8 overexpression inhibits tissue growth through a systemic relay involving Lgr3-positive neurons that connect, via PTTH neurons, to the prothoracic gland where ecdysone is produced (Colombani et al., 2015). Second, ecdysone was shown to promote growth of imaginal tissues (Delanoue et al., 2010;Dye et al., 2017;Herboso et al., 2015;Jaszczak et al., 2015). Third, feeding larvae with ecdysone prevents IOC (Parker and Shingleton, 2011).

### A new pathway controlling Dilp8-dependent growth coordination

Two independent pathways have previously been shown to trigger *dilp8* expression in response to modifications of the growth status. In neoplastic growth conditions, the JNK pathway is a strong inducer of Dilp8 and inhibits the larval-pupal transition in response to developing tumors (Colombani et al., 2012). Additionally, the transcriptional effector of Hippo signaling, Yorkie/Scalloped, modulates the Dilp8 promoter to couple normal growth and *dilp8* expression (Boone et al., 2016). We show here that none of these pathways is responsible for *dilp8* induction in response to growth inhibition in minute tissues. Instead, an independent pathway triggered by Xrp1 is potently activated in these conditions, leading to a strong induction of *dilp8*.

Therefore, different types of growth perturbations (minor growth defects due to developmental variability, neoplastic growth, irradiation, growth inhibition in minute conditions) converge on the regulation of Dilp8 to trigger systemic responses, but use selective, independent pathways to do so. This potentially allows organisms to finely tune local and systemic responses according to the type of growth impairment.

### Xrp1 is a growth inhibitor

Xrp1 was previously involved in genotoxic stress (irradiation) response, and the maintenance of genome integrity downstream of p53 (Akdemir et al., 2007) and during P-element dysgenesis (Francis et al., 2016). Our results uncover a new function for Xrp1 as a growth regulator. The knock-down of *xrp1* in minute discs revealed that it is required for non-autonomous growth inhibition through Dilp8 induction. However, when Rp genes are silenced, removal of Xrp1 is not sufficient to rescue growth autonomously. By contrast, *xrp1* overexpression inhibits tissue growth both autonomously and non-autonomously, and only the non-autonomous response relies on Dilp8. Our study of IOC sheds new light on the biology of Xrp1, demonstrating that Xrp1 carries two independent functions in growth control. On one hand, Xrp1 autonomously inhibits tissue growth. This is in line with previous results showing that Xrp1 is sufficient to inhibit cell proliferation in S2 cells (Akdemir et al., 2007), in eye discs (Tsurui-Nishimura et al., 2013), and with the induction of apoptosis we observed upon Xrp1 overexpression in the wing pouch. On the other hand, activation of Xrp1 signaling in minute discs or upon overexpression remotely inhibits tissue growth in a Dilp8-dependent manner.

### Rps12 as a sensor for ribosomal protein function

Whether the Xrp1 pathway is activated in minute discs by a general reduction of protein translation and/or growth potential, or by a specific pathway such as the ribosomal stress response, is unclear. Our finding that RpS12 is required for IOC and Dilp8 upregulation in minute discs brings insights into this mechanism. RpS12 is one of the last ribosomal proteins assembled into the small ribosomal subunit and, in contrast to most *Drosophila* Rp-encoding genes, *rps12* is not haplo-insufficient. Kale et al. recently established that RpS12 is required to trigger minute cell competition, and that the relative levels of RpS12 define cell competitiveness. Moreover, recent data from Baker’s group indicates that RpS12 is required for *xrp1* expression in minute clones (Lee et al., 2018). We bring here the remarkable finding that removing RpS12, while inducing autonomous growth inhibition by itself, rescues the Xrp1/Dilp8 non-autonomous response in the context of already minute discs. This is indeed strong evidence that Rps12 is itself required for the cells to respond to a defect in Rp protein function. We therefore propose a model whereby RpS12 acts as an upstream signal for IOC upon dis-function of ribosomal proteins (RpS3 or RpL7 in this study). RpS12 is required for Xrp1 activation and *dilp8* upregulation, ultimately leading to systemic response and IOC (Figure 4E). This raises the interesting possibility that ribosomal protein function is used by cells and tissue as a proxy for their growing status and, as such, participates in inter-organ growth coordination mechanisms.

### Xrp1, cell competition and size adjustment

A recent study reports that Xrp1 is required in minute clones to induce cell competition by surrounding wild-type cells (Lee et al., 2018). Closer analysis revealed that Xrp1 mediates most of the changes in gene expression and translation inhibition observed in minute cells. Lee et al. also report a mild transcriptional increase of *xrp1* in heterozygous minute cells, independently of p53. This suggests similarities for the induction of Xrp1 signaling in the context of cell competition and in inter-organ growth coordination. According to Lee et al., Xrp1 is required in minute clones for autonomous growth inhibition. This differs from our study, where Xrp1 is required in minute tissues to trigger the non-autonomous inhibition of growth in other discs, but not within the minute domain itself (see our Figure 2C). This difference might reflect the biological contexts of minute cells in the two experimental setups. Indeed, in minute clones, cells are subjected to direct interactions with surrounding *wt* cells, which induce their elimination by apoptosis. By contrast, in the present study we generate minute conditions in full disc domains, in which case cell competition is limited. Finally, our findings showing that Xrp1 triggers systemic responses raise the possibility that systemic signals could also contribute to local cell competition.

## EXPERIMENTAL PROCEDURES

### Drosophila strains

The following strains were provided by the Bloomington Drosophila Stock Center (BDSC): *nub-Gal4* (38418), *pdm2-Gal4* (49828), *UAS-GFP^RNAi^* (35786), *UAS-CAT* (24621), *UAS-SOD^12.1^* (33605), *xrp1-lacZ* (11569). The following strains were provided by the Vienna Drosophila RNAi Center (VDRC): *UAS-RpS3^RNAi^* (37741), *UAS-RpL7^RNAi^* (21973), *UAS-dilp8^RNAi^* (102604), *UAS-tak1^RNAi^* (101357), *UAS-yki^RNAi^* (104523), *UAS-xrp1^RNAi^* (107860), *UAS-sd^RNAi^* (101497), *UAS-p53^RNAi^* (38235), *UAS-nrf2^RNAi^* (108127), *UAS-mafS^RNAi^* (109303), *UAS-p38^RNAi^* (102484), *UAS-GCN2^RNAi^* (103976), *UAS-PERK^RNAi^* (110278), *UAS-rpS12^RNAi^* (109381). *UAS-dilp8* (Colombani et al., 2012) and *UAS-EcR^RNAi^* (Roignant et al., 2003) were as described. The *elav-GAL80* was kindly provided by Alex Gould (The Francis Crick Institute). Flies were reared and experiments were performed on fly food containing, per liter: 34g inactivated yeast powder, 83g corn flour, 10g agar and 3.5g Moldex. Experiments were done at 25°C.

### Immunohistochemistry

Tissues were dissected in 1X phosphate-buffered saline (PBS) at the indicated hours after egg deposition (AED), fixed in 4% formaldehyde (Sigma) in PBS for 20 minutes at room temperature, washed in PBS containing 0.1% Triton-X-100 (PBT), blocked in PBT containing 10% fetal bovine serum and incubated overnight with primary antibodies at 4 °C. The next day, tissues were washed, blocked again and incubated with secondary antibodies at 1/400 dilution for 2 hours at room temperature. After washing, tissues were mounted in Vectashield containing DAPI for staining of DNA (Vector Labs). Fluorescence images were acquired using a Leica SP5 DS confocal microscope and processed using Adobe Photoshop CS5 or Image J. For Dilp8 stainings, z stacks of 10μm (0,29μm/slice) were acquired starting on the apical side on the wing discs and presented as maximal projections (Image J).

The following primary antibodies were used: mouse anti-Wingless (Wg; concentrated 4D4 from DSHB; 1/200), rabbit anti-cleaved Caspase 3 (Casp3*; Cell Signaling; 1/100), chicken anti-beta-galactosidase (β-Gal; GeneTek; 1/1000), rat anti-Dilp8 (Colombani et al., 2012; 1/500). The following secondary antibodies were used: Alexa Fluor 488 goat anti-mouse, Alexa Fluor 546 goat anti-rabbit, Alexa Fluor 546 goat anti-rat, Alexa Fluor 488 goat anti-chicken.

### Pupariation curves

4-hours egg layings were collected on agar plates with yeast. L1 larvae were collected 24 hours AED and reared in tubes (forty larvae each) containing standard food (see above). The number of larvae that had pupariated was scored at the indicated time points AED.

### Quantitative RT-PCR

Larvae were collected at the indicated number of hours AED. Whole larvae or dissected wing discs were frozen in liquid nitrogen. Total RNA was extracted using a QIAGEN RNeasy Lipid Tissue Mini Kit (for whole larvae) or a QIAGEN RNeasy Micro Kit (for dissected wing discs) according to the manufacturer’s protocol. RNA samples (2-3μg per reaction) were treated with DNase and reverse-transcribed using SuperScript II reverse transcriptase (Invitrogen), and the generated cDNAs were used for real-time PCR (StepOne Plus, Applied Biosystems) using PowerSYBRGreen PCR mastermix (Applied Biosystems). Samples were normalized to the levels of the ribosomal protein rp49 transcript levels and fold changes were calculated using the ΔΔCt method. Three separate biological samples were collected for each experiment and triplicate measurements were performed. The following primers were used: rp49-sense: 5’-CTT CAT CCG CCA CCA GTC-3’, rp49-antisense: 5’-CGA CGC ACT CTG TTG TCG-3’, dilp8-sense: 5’-CGA CAG AAG GTC CAT CGA GT-3’, dilp8-antisense: 5’-GTT TTG CCG GAT CCA AGT C-3’, xrp1-L-sense: 5’-TCA TTG TTT CTT TCT AAC GGT CAA-3’, xrp1-L-antisense: 5’-GGT TGC TGT TGT TTG ATT CG-3’, xrp1-common-sense: 5’-GAC CAC ACC GGA GAT TAT CAA-3’, xrp1-common-antisense: 5’-GCT GGT ACT GGT ACT TGT GGT G-3’.

### Xrp1 cloning

The sequence encoding the long form of Xrp1 (Xrp1-L; amino acids 1-668) was PCR amplified from the BDGP EST cDNA clone SD01985 and cloned into the pENTR/D-TOPO vector using the following primers: sense: 5’-CAC CAT GAT CCA GGA GCC AGC ACG AGT A-3’ and antisense: 5’-TCA GTC CTG CTC CTG CTT AAC GTA AG-3’ (with stop codon). Sequence analysis detected a number of mutations in the SD01985 clone (according to the reference genome assembly for *D. melanogaster*) that were corrected using the QuickChange site-directed mutagenesis kit (Stratagene). The sequence encoding the short form of Xrp1 (Xrp1-S; amino acids 263-668) was PCR amplified from *w^1118^* cDNA and cloned into the pENTR/D-TOPO vector using the following gene-specific primers: sense: 5’-CAC CAT GTT TGC CGA GGA GGA TCT GAT-3’ and antisense 5’-TCA GTC CTG CTC CTG CTT AAC GTA AG-3’ (with stop codon). To generate transgenic lines harboring the xrp1-L and xrp1-S coding sequences under the control of UAS (UAS-xrp1-L and UAS-xrp1-S), the pUASattB-xrp1-S and pUASattB-xrp1-L constructs were injected in the presence of the PhiC31 integrase and inserted into the 51C landing site on the 2nd chromosome.

### Statistics

P values are the result of ANOVA tests provided by GraphPad Prism (^**^ p <0.01 and ^***^ p<0.001).

## ACKNOWLEDGMENTS

We thank T. Pihl, J. Villalba and L. Ruel for help with the experiments; M. Milan, K. Basler, the Bloomington Stock Center and the Vienna Drosophila RNAi Center for fly stocks; K. Basler, D. Lubensky, N. Baker, M. Milan, J-P. Vincent for discussions. This project has received funding from the Association pour la recherche sur le cancer (grant n°PGA120150202355 to P.L.), Inserm, CNRS, the European Research Council (ERC Advanced grant n°268813 to P.L.), the Labex Signalife program (grant ANR-11-LABX-0028-01 to P.L.), the Marie Sklodowska-Curie Actions (fellowship n°657685 to L.B.).

## FIGURES LEGENDS

**Figure S1.**
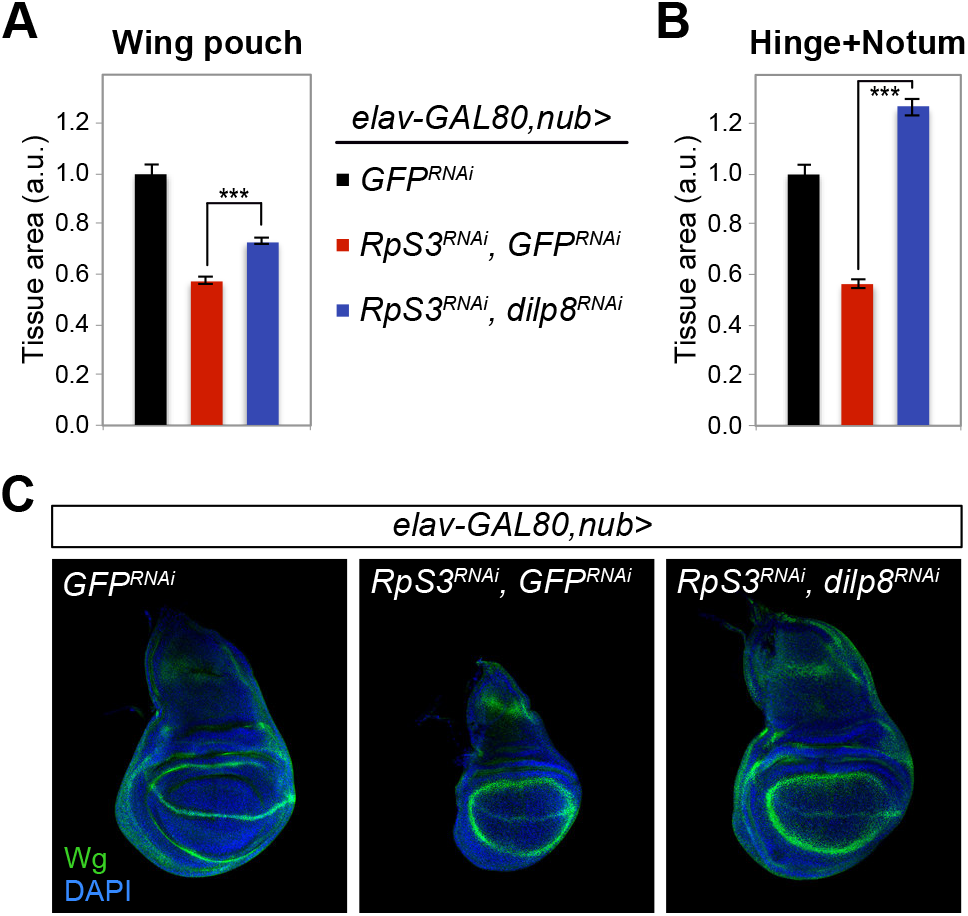
Dilp8 is required for growth coordination in *elav-GAL80, nub>RpS3^RNAi^* wing discs. (A,B) Measurement of wing pouch area (A) and hinge and notum area (B) in control wing discs (*elav-GAL80, nub>GFP^RNAi^*), minute wing discs (*elav-GAL80, nub>RpS3^RNAi^, GFP^RNAi^*), and minute wing discs with the simultaneous knock-down of *dilp8* (*elav-GAL80, nub>RpS3^RNAi^, dilp8^RNAi^*). (C) Representative pictures of wing imaginal discs of the three genotypes stained for Wingless (Wg) illustrate the loss of coordination between the different territories in *elav-GAL80, nub>RpS3^RNAi^, dilp8^RNAi^* wing discs. Experiments were done at 110h AED (n≥20). Data are represented as mean ± SEM (*** p<0.001, ANOVA).

**Figure S2:**
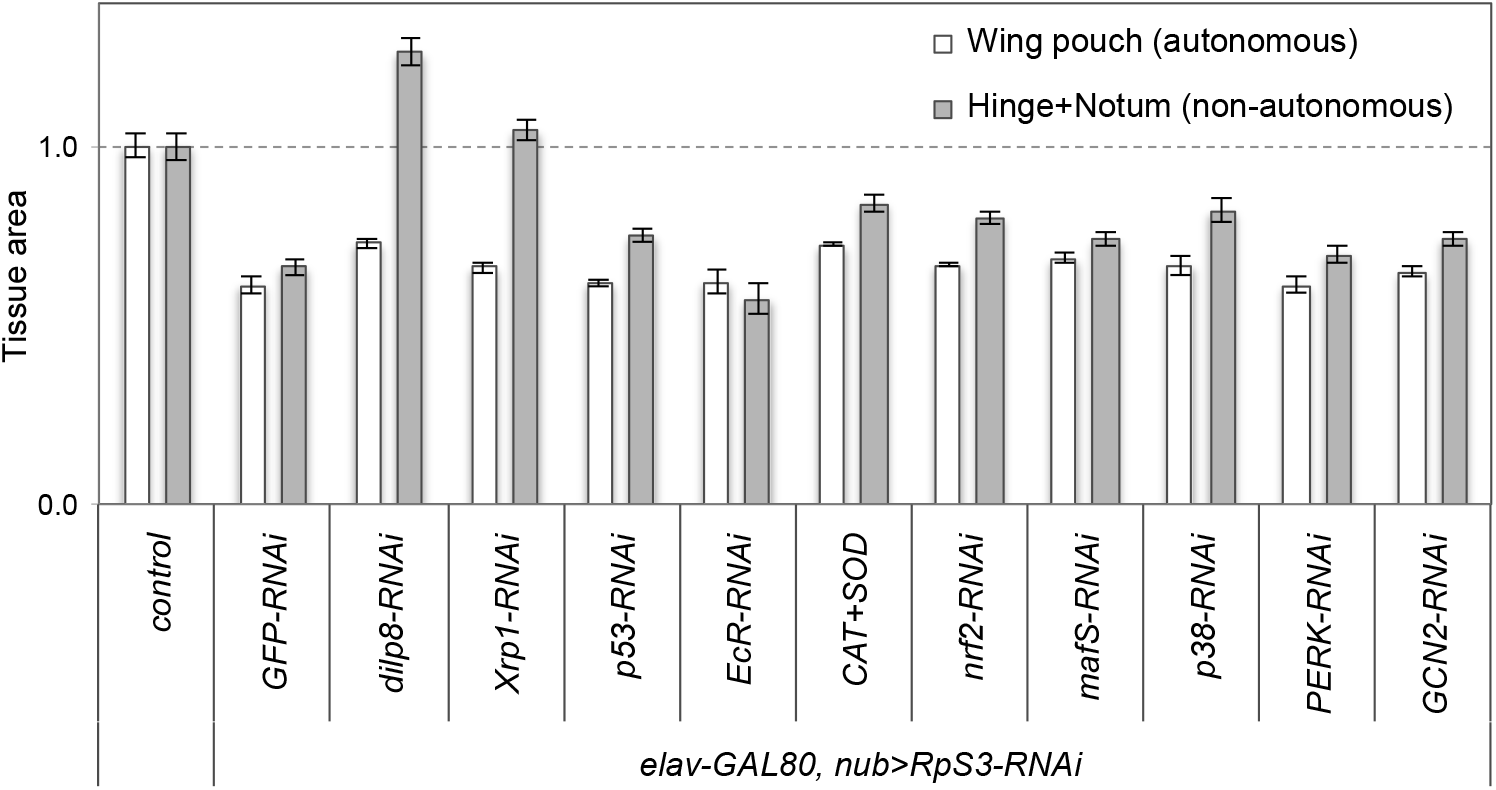
Summary of the different signaling pathways tested for growth coordination. Tissue area, expressed as a ratio to control, of the wing pouch and hinge+notum territories in the indicated genetic conditions. Experiments were done at 110h AED (n≥13). Data are represented as mean ± SEM.

**Figure S3:**
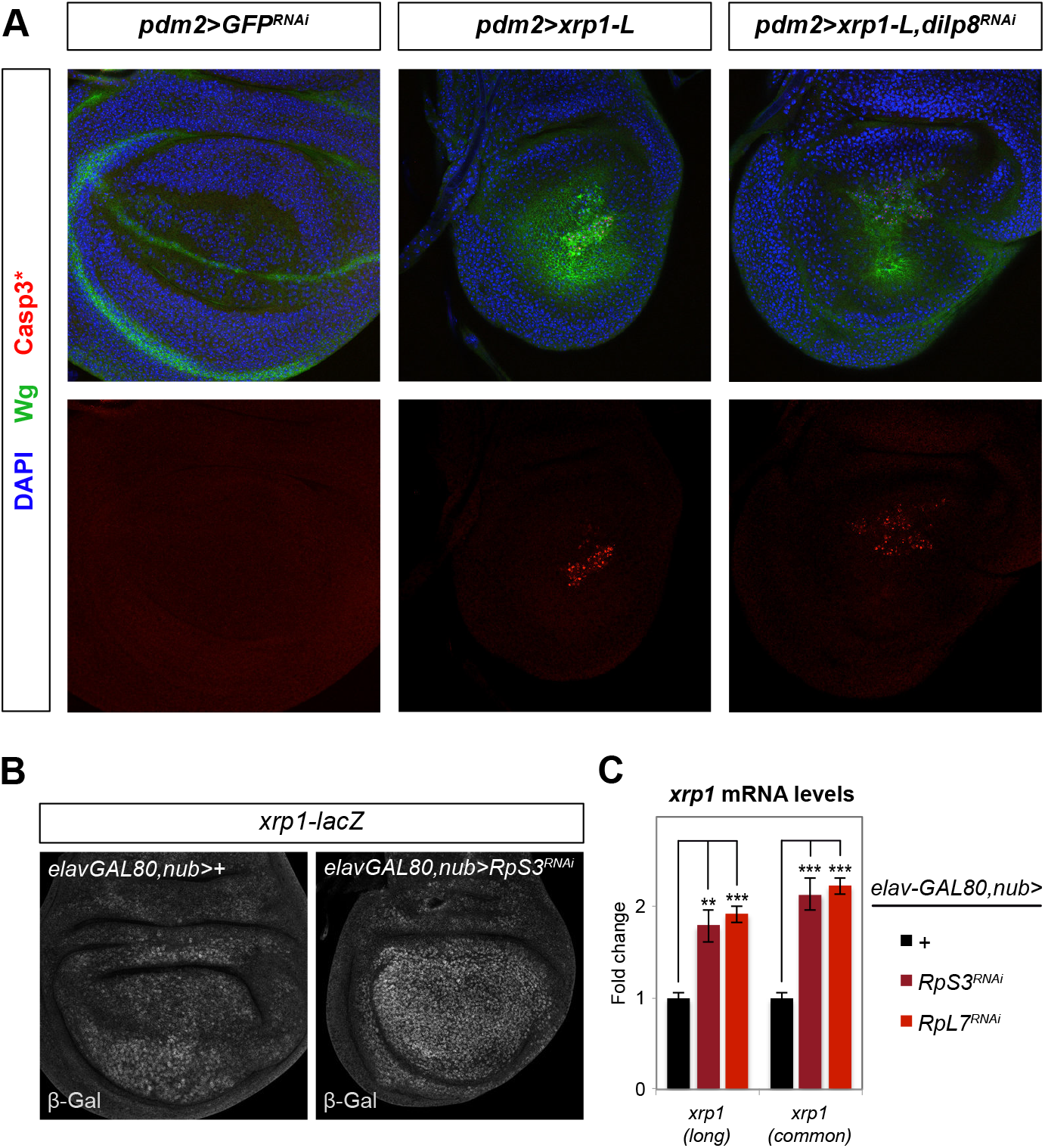
Induction of cell death by Xrp1 and transcriptional regulation of *xrp1*. (A) Xrp1 overexpression induces cell death autonomously in a Dilp8-independent manner. Representative pictures of wing imaginal discs of the different genotypes stained for Wg and activated caspase 3 (Casp3^*^, indicative of cell death). (B) Activity of the *xrp1-lacZ* reporter in control and minute wing pouch. (C) *xrp1* mRNA levels measured by qRT-PCR on dissected wing discs of control and minute discs. Experiments were done at 110h AED. Data are represented as mean ± SEM (^**^ p <0.01 and ^***^ p<0.001, ANOVA).

**Figure S4.**
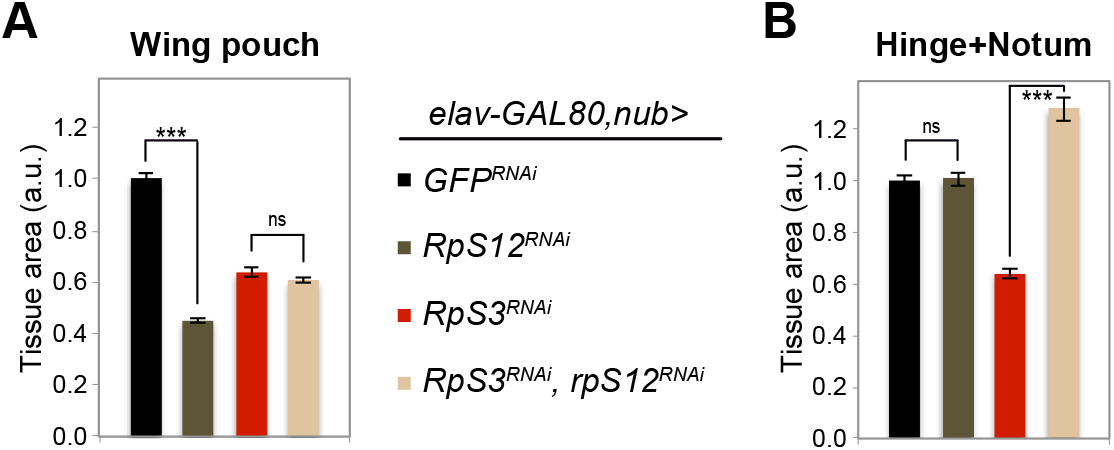
RpS12 is required for growth coordination in *elav-GAL80, nub>RpS3^RNAi^* wing discs. (A-C) Measurement of wing pouch area (A) and hinge and notum area (B) in control wing discs (*elav-GAL80, nub>GFP^RNAi^*), wing discs with knock-down of *rpS12* (*elav-GAL80, nub>rpS12^RNAi^*), minute wing discs (*elav-GAL80, nub>RpS3^RNAi^, GFP^RNAi^*), and minute wing discs with simultaneous knock-down of *rpS12* (*elav-GAL80, nub>RpS3^RNAi^, rpS12^RNAi^*). Experiments were done at 110h AED (n≥30). Data are represented as mean ± SEM (^***^ p<0.001 and ns=not significant, ANOVA).

## REFERENCES

Akdemir, F., Christich, a., Sogame, N., Chapo, J., and Abrams, J.M. (2007). p53 directs focused genomic responses in Drosophila. Oncogene 26, 5184–5193.

Boone, E., Colombani, J., Andersen, D.S., and Léopold, P. (2016). The Hippo signalling pathway coordinates organ growth and limits developmental variability by controlling dilp8 expression. Nat. Commun. 7, 13505.

Colombani, J., Andersen, D.S., and Léopold, P. (2012). Secreted peptide Dilp8 coordinates Drosophila tissue growth with developmental timing. Science 336, 582–585.

Colombani, J., Andersen, D.S., Boulan, L., Boone, E., Romero, N., Virolle, V., Texada, M., and Léopold, P. (2015). Drosophila Lgr3 Couples Organ Growth with Maturation and Ensures Developmental Stability. Curr. Biol. 25.

Delanoue, R., Slaidina, M., and Leopold, P. (2010). The steroid hormone ecdysone controls systemic growth by repressing dMyc function in Drosophila fat cells. Dev Cell 18, 1012–1021.

Demay, Y., Perochon, J., Szuplewski, S., Mignotte, B., and Gaumer, S. (2014). The PERK pathway independently triggers apoptosis and a Rac1/Slpr/JNK/Dilp8 signaling favoring tissue homeostasis in a chronic ER stress Drosophila model. Cell Death Dis. 5, e1452–10.

Dye, N.A., Popović, M., Spannl, S., Etournay, R., Kainmüller, D., Ghosh, S., Myers, E.W., Jülicher, F., and Eaton, S. (2017). Cell dynamics underlying oriented growth of the Drosophila wing imaginal disc. Development 144, 4406–4421.

Francis, M.J., Roche, S., Cho, M.J., Beall, E., Min, B., Panganiban, R.P., and Rio, D.C. (2016). *Drosophila* IRBP bZIP heterodimer binds P-element DNA and affects hybrid dysgenesis. Proc. Natl. Acad. Sci. 113, 13003–13008.

Garelli, A., Gontijo, A.M., Miguela, V., Caparros, E., and Dominguez, M. (2012). Imaginal discs secrete insulin-like peptide 8 to mediate plasticity of growth and maturation. Science 336, 579–582.

Gruenewald, C., Botella, J. a., Bayersdorfer, F., Navarro, J. a., and Schneuwly, S. (2009). Hyperoxia-induced neurodegeneration as a tool to identify neuroprotective genes in Drosophila melanogaster. Free Radic. Biol. Med. 46, 1668–1676.

Herboso, L., Oliveira, M.M., Talamillo, A., Pérez, C., González, M., Martín, D., Sutherland, J.D., Shingleton, A.W., Mirth, C.K., and Barrio, R. (2015). Ecdysone promotes growth of imaginal discs through the regulation of Thor in D. melanogaster. Sci. Rep. 5, 12383.

Jaszczak, J.S., and Halme, A. (2016). Arrested development: coordinating regeneration with development and growth in Drosophila melanogaster. Curr. Opin. Genet. Dev. 40, 87–94.

Jaszczak, J.S., Wolpe, J.B., Dao, A.Q., and Halme, A. (2015). Nitric oxide synthase regulates growth coordination during Drosophila melanogaster imaginal disc regeneration. Genetics.

Kale, A., Ji, Z., Kiparaki, M., Blanco, J., Rimesso, G., Flibotte, S., and Baker, N.E. (2018). Ribosomal Protein S12e Has a Distinct Function in Cell Competition. Dev. Cell 44, 42–55.e4.

Katsuyama, T., Comoglio, F., Seimiya, M., Cabuy, E., and Paro, R. (2015). During Drosophila disc regeneration, JAK/STAT coordinates cell proliferation with Dilp8-mediated developmental delay. Proc. Natl. Acad. Sci. U. S. A. 112, E2327–36.

Kucinski, I., Dinan, M., Kolahgar, G., and Piddini, E. (2017). Chronic activation of JNK JAK/STAT and oxidative stress signalling causes the loser cell status. Nat. Commun.

Lee, C.-H., Rimesso, G., Reynolds, D.M., Cai, J., and Baker, N.E. (2016). Whole-Genome Sequencing and iPLEX MassARRAY Genotyping Map an EMS-induced Mutation Affecting Cell Competition in Drosophila melanogaster. G3 6, 3207–3217.

Lee, C.-H., Kiparaki, M., Blanco, J., Folgado, V., Ji, Z., Kumar, A., Rimesso, G., and Baker, N.E. (2018). A Regulatory Response to Ribosomal Protein Mutations Controls Translation, Growth, and Cell Competition. Dev. Cell 46, 1–14.

Parker, N.F., and Shingleton, A.W. (2011). The coordination of growth among Drosophila organs in response to localized growth-perturbation. Dev. Biol. 357, 318–325.

Roignant, J.Y., Carré, C., Mugat, B., Szymczak, D., Lepesant, J.A., and Antoniewski, C. (2003). Absence of transitive and systemic pathways allows cell-specific and isoform-specific RNAi in Drosophila. RNA.

Roselló-Díez, A., Madisen, L., Bastide, S., Zeng, H., and Joyner, A.L. (2018). Cell-nonautonomous local and systemic responses to cell arrest enable long-bone catch-up growth in developing mice. PLoS Biol. 16, e2005086.

Tsurui-nishimura, N., Nguyen, T.Q., Katsuyama, T., Minami, T., Furuhashi, H., Oshima, Y., and Kurata, S. (2013). Ectopic Antenna Induction by Overexpression of CG17836/Xrp1 Encoding an AT-Hook DNA Binding Motif Protein in Drosophila. Biosci. Biotechnol. Biochem. 77, 339–344.

Zhang, C., Robinson, B.S., Xu, W., Yang, L., Yao, B., Zhao, H., Byun, P.K., Jin, P., Veraksa, A., and Moberg, K.H. (2015). The Ecdysone Receptor Coactivator Taiman Links Yorkie to Transcriptional Control of Germline Stem Cell Factors in Somatic Tissue. Dev. Cell 34, 168–180.

